# 1-deoxysphingolipid biosynthesis compromises anchorage-independent growth and plasma membrane endocytosis in cancer cells

**DOI:** 10.1101/2022.01.19.476986

**Authors:** Thekla Cordes, Ramya S. Kuna, Grace H. McGregor, Sanika V. Khare, Jivani Gengatharan, Thangaselvam Muthusamy, Christian M. Metallo

## Abstract

Serine palmitoyltransferase (SPT) predominantly incorporates serine and fatty acyl-CoAs into diverse sphingolipids that serve as structural components of membranes and signaling molecules within or amongst cells. However, SPT also uses alanine as a substrate in the contexts of low serine availability, alanine accumulation, or disease-causing mutations in hereditary sensory neuropathy type I (HSAN1), resulting in the synthesis and accumulation of 1-deoxysphingolipids. These species promote cytotoxicity in neurons and impact diverse cellular phenotypes, including suppression of anchorage-independent cancer cell growth. While altered serine and alanine can promote 1-deoxysphingolipid synthesis, they impact numerous other metabolic pathways important for cancer cells. Here we combined isotope tracing, quantitative metabolomics, and functional studies to better understand the mechanistic drivers of 1-deoxysphingolipid toxicity in cancer cells. Both alanine treatment and *SPTLC1^C133W^* expression induce 1-deoxy(dihydro)ceramide synthesis and accumulation but fail to broadly impact intermediary metabolism, abundances of other lipids, or growth of adherent cells. However, spheroid culture and soft agar colony formation were compromised when endogenous 1-deoxysphingolipid synthesis was induced via *SPTLC1^C133W^*expression. Consistent with these impacts on anchorage-independent cell growth, we observed that 1-deoxysphingolipid synthesis reduced plasma membrane endocytosis. These results highlight a potential role for SPT promiscuity in linking altered amino acid metabolism to plasma membrane endocytosis.

## INTRODUCTION

Sphingolipids (SLs) are bioactive molecules and structural components of the cell membrane influencing essential cellular processes (1). Serine palmitoyltransferase (SPT) catalyzes the condensation of palmitate and serine in *de novo* biosynthesis of SLs but can also produce non-canonical, cytotoxic 1-deoxysphingolipids (deoxySLs) when using alanine or glycine as a substrate (2-4). Alterations in serine and alanine levels impact substrate availability for the promiscuous SPT reaction driving the accumulation of non-canonical SLs (5,6). DeoxySLs differ from canonical SL in that they lack the C1-hydroxyl group of sphinganine, preventing further modification or degradation via canonical SL pathways (3). DeoxySLs are cytotoxic to cells, though specific mechanisms remain unclear (7-9). Accumulation of deoxySLs in the contexts of extracellular matrix detachment and/or serine deprivation compromises anchorage-independent growth and the establishment of xenograft tumors (10). However, serine and glycine restriction impact numerous downstream pathways, including nucleotide synthesis/one carbon metabolism (11-14), NAD(P)H (15-17), and lipid metabolism (10,18). On the other hand, exogenous addition of deoxySLs impacts diverse cellular processes, including mitochondrial function (8), autophagy (19), migration (20), sphingosine kinase 1 proteolysis (21,22), and protein folding (23), but this approach does not recapitulate the location and trafficking of deoxySLs actively synthesized on the endoplasmic reticulum (ER). We therefore aimed to determine deoxySL specific toxic effects independent of the broader metabolic impacts of serine and glycine restriction.

Alanine is abundant in the plasma and accumulates in diseases associated with mitochondrial dysfunction (24), metabolic syndrome (25,26), or anoxia (23). Transaminases expressed in cancer cells readily metabolize alanine and alpha-ketoglutarate to pyruvate and glutamate, but a reduction in pyruvate oxidation will cause alanine accumulation and subsequent conversion to deoxySA (10). Alanine supplementation therefore offers a means of driving deoxySL accumulation without restricting serine. Further, deoxySLs accumulate in the context of point mutations in *SPTLC1* or *SPTLC2*, which encode subunits of the SPT complex, occur in patients with hereditary sensory neuropathy type I (HSAN1), and increase the complex’s affinity for alanine (2,4,27,28). Ectopic expression of such *SPTLC1* variants effectively drives the endogenous accumulation of deoxySL in cells (27,29–31) offering an additional means of probing deoxySL-specific cellular effects.

Here we investigate how deoxySL biosynthesis alters membrane lipid metabolism and cellular processes, comparing 2D cell growth to anchorage-independent conditions. By combining high-resolution mass spectrometry, fluorescence imaging, and anchorage-independent growth studies, we highlight key aspects of deoxySLs on cancer cell metabolism and growth. These results demonstrate the impact of deoxySL on soft agar colony growth and endocytosis at the plasma membrane, illustrating key functional impacts relevant for cancer and neuropathy.

## RESULTS

### Alanine promotes deoxySL synthesis and compromises early spheroid formation

To better understand how cytotoxic deoxySLs impact cell function, we altered alanine availability to drive endogenous deoxySL biosynthesis in 2D cultures. Alanine is highly synthesized in proliferating mammalian cells cultured *in vitro* but absent from most media. Supplementation of 0.5 - 1 mM alanine increased deoxy-dihydroceramides (deoxyDHCer) and deoxy-ceramides (deoxyCer) pools by ∼50%. At the same time, levels of canonical dihydroceramides (DHCer) and ceramides (Cer) were not significantly affected (Fig. 1A). We also observed that UK5099, an inhibitor of the mitochondrial pyruvate carrier (MPC), decreased alanine levels by promoting its oxidation (Figs. 1B, S1A). In turn, modulation of MPC and alanine levels had a strong impact on free deoxySA m18:0 in HCT116 cells (Fig. 1C). Further, numerous deoxyDHCer and deoxyCer species increased in response to alanine while UK5099 buffered their accumulation (Figs. 1D, S1B). These data highlight the direct impact of alanine and MPC function on deoxySL accumulation. Notably, exogenous alanine did not impact cell growth in 2D adherent culture, despite increasing non-canonical deoxySL species (Fig. 1E). We recapitulated our findings using A549 non-small lung cancer cells, further indicating that different cell types can accumulate deoxySL without exhibiting cytotoxicity (Figs. S1C, D).

**Figure 1:**
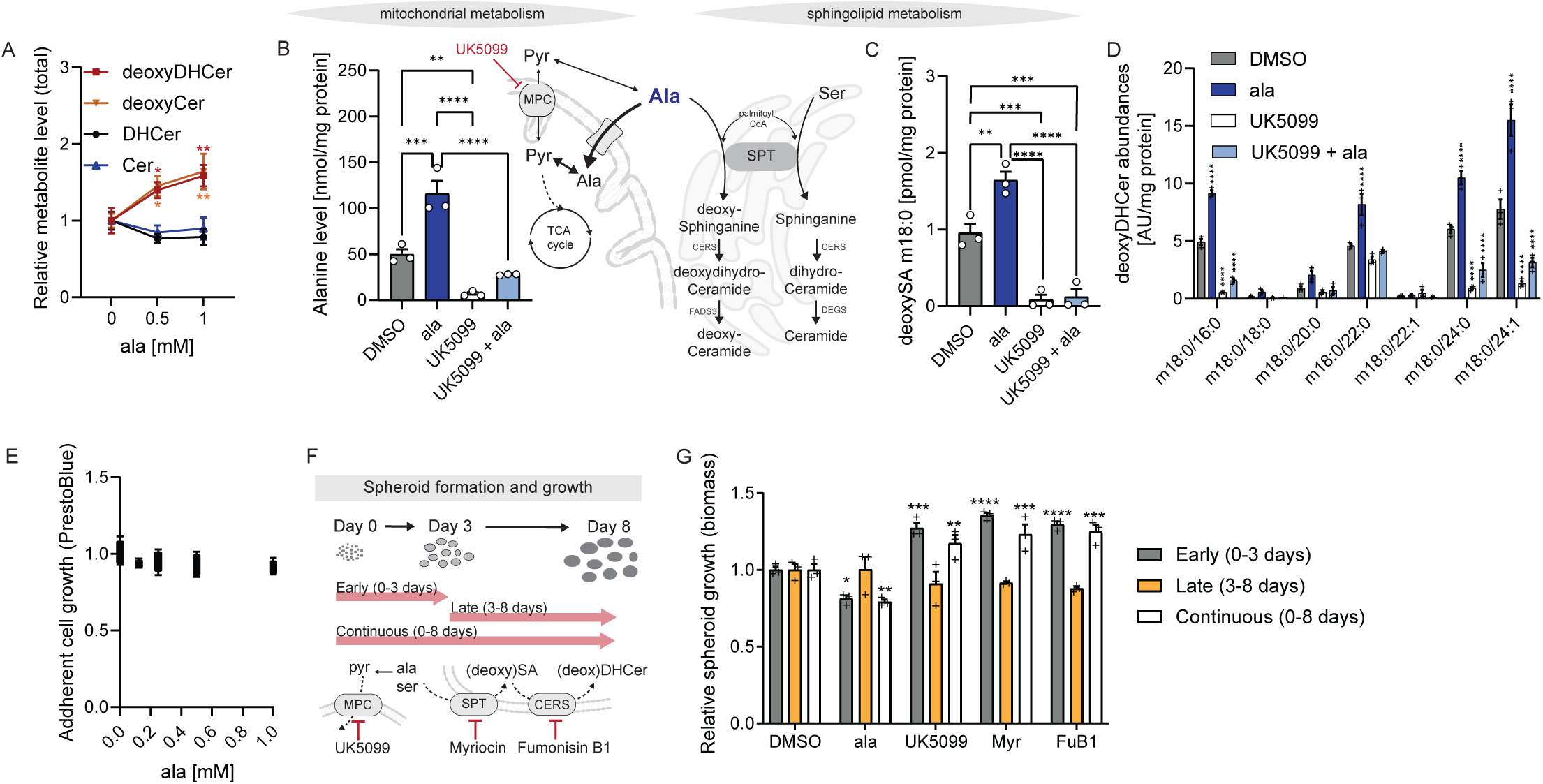
Alanine promotes deoxySL synthesis and compromises early spheroid formation. A. Total deoxyDHCer, deoxyCer, DHCer, and Cer levels in HCT116 cells cultured with increasing alanine for 3 days relative to 0 mM alanine. B. Alanine levels in HCT116 cells cultured for 3 days with 1 mM alanine and 5 µM UK5099. Schematic depicting alanine involved in mitochondrial and sphingolipid metabolism. SPT: serine palmitoyltransferase; MPC: mitochondrial pyruvate carrier C. DeoxySA m18:0 levels in HCT116 cells cultured for 3 days with 1 mM alanine and 5 µM UK5099. D. Levels of deoxyDHCer species in HCT116 cells cultured for 3 days with 1mM alanine and 5 µM UK5099. E. Adherent cell growth of HCT116 cells cultured with increasing alanine concentrations for 3 days. F. Schematic depicting spheroid formation assay with inhibitors targeting MPC (UK5099), SPT (myriocin), and ceramide synthase (CERS, fumonisin B1). G. HCT116 spheroid growth with DMSO, 1 mM alanine, 5 µM UK5099, 10 nM myriocin, 10 µM fumonisin B1 (FeB1) for 3 to 8 days as indicated in the figure. Cells were grown in adherent (A-E) or spheroid cultures (G). Data are presented as means ± s.e.m. (A-D, G) with three cellular replicates or box (25^th^ to 75^th^ percentile with median line) and whiskers (min. to max. values) (E) with five cellular replicates. One-way ANOVA (B, C) or two-way ANOVA (A, D, G). Significance in A is compared to 0mM alanine. Significance for D and G is compared to DMSO. Results are depicted from one representative experiment and each experiment was repeated independently three times with similar results. * *P* < 0.05, ** P < 0.01, *** *P* < 0.001, **** *P* < 0.0001.

In contrast to adherent culture models, we have previously demonstrated that alanine supplementation and deoxySL accumulation specifically compromises spheroid growth (10). Spheroid growth requires cells to initially survive upon trypsinization and sub-culture into anchorage-independent conditions; this process has a profound influence on glycans and downstream metabolic processes important for rebuilding these membrane structures (32). Subsequently, cells in spheroid culture must sustain proliferation in larger 3D structures which establish nutrient and hypoxic gradients and associated imbalances in redox pathways (33–35). To better understand how alanine and deoxySL metabolism impact spheroid growth we plated cells in low attachment plates and modulated deoxySL synthesis by supplementing alanine or inhibiting the MPC (UK5099), the SPT complex (myriocin), or ceramide synthases (CERS) (fumonisin B1). We added treatments at early (days 0-3) or late (days 3-8) spheroid formation phase and compared our results to spheroid treated continuously (days 0-8) as previously described (10) (Fig. 1F). Intriguingly, we observed an impact on spheroid biomass only when cells were treated in the early phase (i.e. Early and Continuous), while treatment during later stages of spheroid growth had no effect (Fig. 1G). Remarkably, these changes were recapitulated by all biochemical treatments, including alanine, UK5099, myriocin, and fumonsin B1. Presumably the 3-8 day “late” treatments drive similar changes in sphingoid bases to those confirmed after 24 hour and 8 day continuous treatments (10); however, these data highlight that alanine and other modulators of deoxySL accumulation compromise growth during the early phase of spheroid formation rather than modulating growth/survival in later-stage, larger spheroid structures. Notably, each of these compounds also impacts other pathways (e.g. canonical sphingolipids, TCA and amino acid metabolism), limiting our ability to confirm promiscuity-linked mechanism using such treatments (and rescues).

### Impact of endogenous deoxySL synthesis on metabolic fluxes

To better elucidate the biochemical and functional impacts of endogenously synthesized deoxySLs independent of alanine supplementation, serine restriction, or canonical sphingolipid synthesis, we established a controllable means of accumulating deoxySLs in isogenic cells. Indeed, ectopic expression of mutant SPTLC1 protein variants or fusion proteins robustly drives deoxySL accumulation in cultured cells (27,29,30). Here, we expressed doxycycline-inducible constructs containing either wild-type (grey: *SPTLC1^WT^*) or mutant (red: *SPTLC1^C113W^*) *SPTLC1* in HCT116 cells via lentiviral delivery. SPTLC1^C113W^ has an increased affinity for alanine, driving the synthesis of deoxySLs compared to SPTLC1^WT^ (Fig. 2A), as observed in patients with HSAN1 (2,4,27). We first confirmed that SPTLC1 protein variants were consistently expressed in cells (Fig. 2B). As expected, levels of deoxyDHCer and deoxyCer accumulated significantly in doxycycline-treated cells expressing *SPTLC1^C133W^*compared to those infected with the *SPTLC1^WT^* construct, while levels of DHCer and Cer were less affected (Fig. 2C). Further, all deoxySL species detected, including free deoxySA (Fig. S2A), deoxyDHCer (Fig. 2D), and deoxyCer (Fig. 2E), were robustly increased in *SPTLC1^C133W^* compared to *SPTLC1^WT^*expressing cells. On the other hand, levels of canonical DHCer and Cer species were less impacted in both polyclonal cell lines (Figs. S2B, S2C).

**Figure 2:**
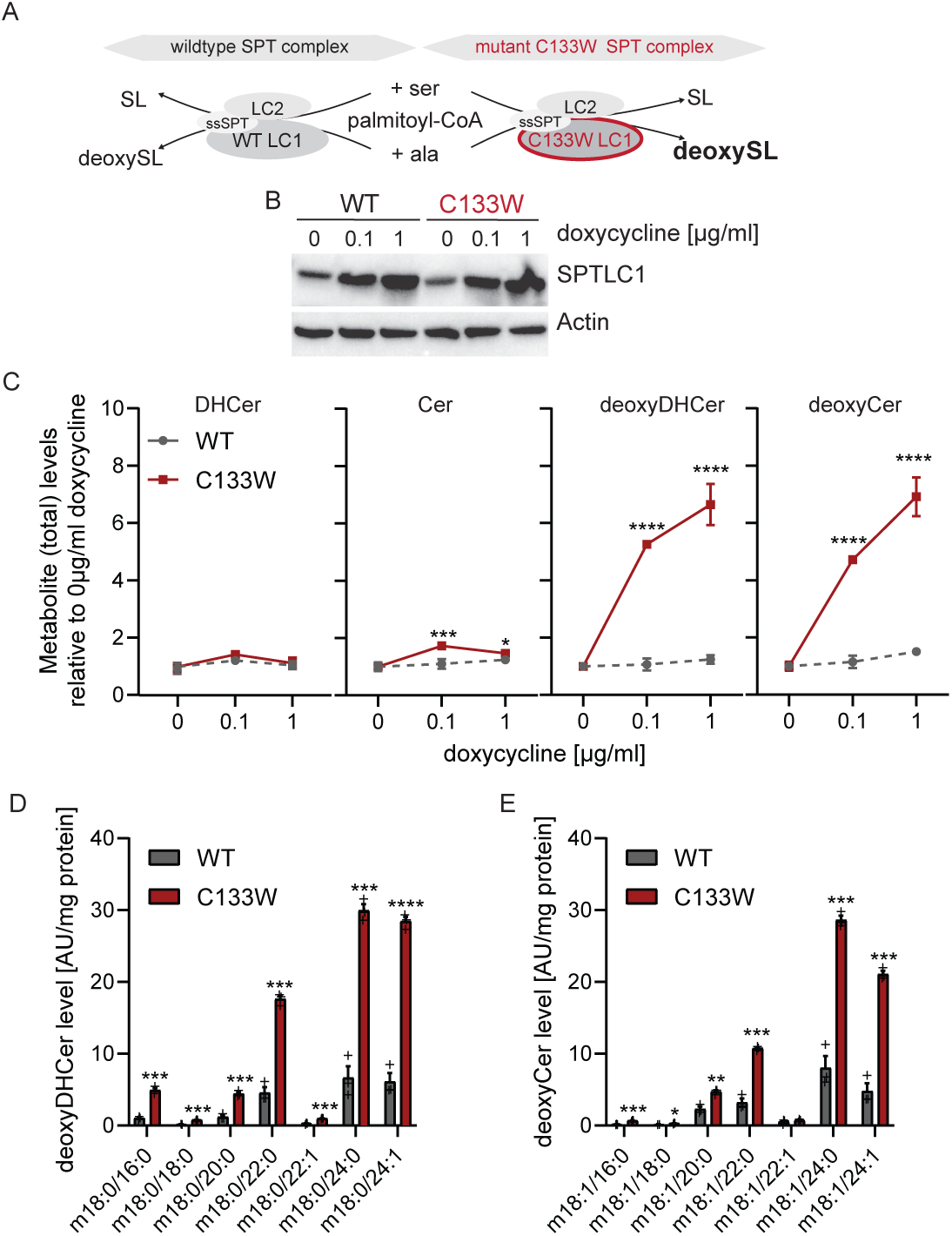
Mutant *SPTLC1*^C133W^ drives non-canonical deoxySL biosynthesis. A. Schematic depicting serine palmitoyl transferase (SPT) promiscuity catalyzing sphingolipid (SL) species from serine or deoxy-sphingolipids (deoxySL) from alanine. Mutant *SPTLC1^C133W^*(C133W, red) has an increased affinity for alanine as substrate compared to *SPTLC1^WT^* (WT, grey) driving deoxySL synthesis. B. Immunoblotting of SPTLC1 and actin in SPT expressing HCT116 cell lines with doxycycline for 4 days. C. Total DHCer, Cer, deoxyDHCer, and deoxyCer levels in HCT116 cells expressing *SPTLC1^WT^* or *SPTLC1^C133W^* cultured for 5 days with increasing doxycycline concentrations. D. Levels of deoxyDHCer species in engineered SPT HCT116 cells cultured for 5 days with 0.1 µg/ml doxycycline. E. Levels of deoxyCer species in engineered SPT HCT116 cells cultured for 5 days with 0.1 µg/ml doxycycline. Cells were grown in adherent cultures and data are presented as means ± s.e.m. with three cellular replicates. Two-way ANOVA compared to 0 µg/ml doxycycline (C) or Student’s t-test (D, E). Experiments were repeated two (B) or three (C-E) times with similar results. \**P* < 0.05, ** *P* < 0.01, *** *P* < 0.001, **** *P* < 0.0001.

To further understand the turnover and metabolism of sphingolipid intermediates in the context of deoxySL synthesis we cultured cells in the presence of [U-^13^C16]palmitate and quantified ^13^C incorporation into sphingoid bases and ceramide species using high-resolution mass spectrometry (Fig. 3A). Although this approach is limited by the non-physiological concentration and localization of the administered [U-^13^C16]palmitate tracer, simultaneous quantitation of SL and deoxySL labeling kinetics is informative of the relative flux through each pathway. Focusing on the more abundant, canonical ceramide pools (i.e., SA d18:0, DHCer d18:0/24:0, and Cer d18:1/24:0) that achieved either steady-state isotopic labeling or >20% turnover, we observed that *SPTLC1^C133W^*cells exhibited reduced labeling kinetics compared to *SPTLC1^WT^*cells (Fig. 3B). Since ceramide pools (including 24:0 n-acyl species) were generally not altered by expression of *SPTLC1* constructs (Figs. S3A, B), these results suggest that *SPTLC1^C133W^*expression or deoxySL accumulation modestly reduced d18:0/24:0 DHCer biosynthesis, but the larger ceramide metabolic network mitigates this impact at this level of *SPTLC1^C133W^* expression (Fig. 2A). On the other hand, turnover of the lower abundance deoxySA m18:0, deoxyDHCer m18:0/24:0, and deoxyCer m18:1/24:0 pools was significantly lower than that observed for canonical sphingolipids (Fig. 3C), which highlights that biosynthetic flux of deoxySL species generally occurs at much slower rates compared to the production of canonical ceramides in *SPTLC1^C133W^*-expressing cells. Similar trends were observed for 24:1 n-acyl (Figs. S3A, B, C, D) as well as 16:0 n-acyl (Figs. S3A, B, E, F) ceramides, in the latter case labeling on both sphingoid bases and n-acyl chains was evident from M+32 isotopologues (Figs. S3G, H). However, labeling kinetics were not as consistently impacted, suggesting that deoxySL synthesis and accumulation have diverse impacts on downstream sphingolipid metabolism depending on the specific molecular species. To determine how deoxySL accumulation influenced other metabolic pathways we quantified broader labeling and flux changes occurring in *SPTLC1^WT^* and *SPTLC1^C113W^*-expressing cells. For example, palmitate oxidation and enrichment of citrate via acetyl-coenzyme A and citrate synthase were unaltered in both cell lines (Fig. 3D). Isotope enrichment from [U-^13^C_6_]glucose was similarly unchanged when deoxySL synthesis was active (Fig. S3I). In addition, doxycycline-treatment and deoxySL accumulation over 6 days failed to impact mitochondrial respiration measured by Seahorse (Fig. S3J). These results suggest that endogenous deoxySL synthesis is not sufficient to significantly alter central carbon metabolism or mitochondrial respiration. Indeed, adherent growth was not impacted by doxycycline-induced expression with increased deoxySL biosynthesis by expression of *SPTLC1^C133W^*, suggesting that cells can accommodate a certain level of *de novo* synthesized deoxySLs (or canonical SLs) without experiencing deleterious effects on growth (Fig. 3E). Consistent with prior results (10) and those presented above (Fig. 1G), we observed a slight but significant reduction in spheroid growth in *SPTLC1^C133W^*compared to *SPTLC1^WT^* HCT116 cells suggesting that the stress from initiating non-adherent growth sensitizes cells to deoxySL accumulation induced toxicity (Fig. 3F).

**Figure 3:**
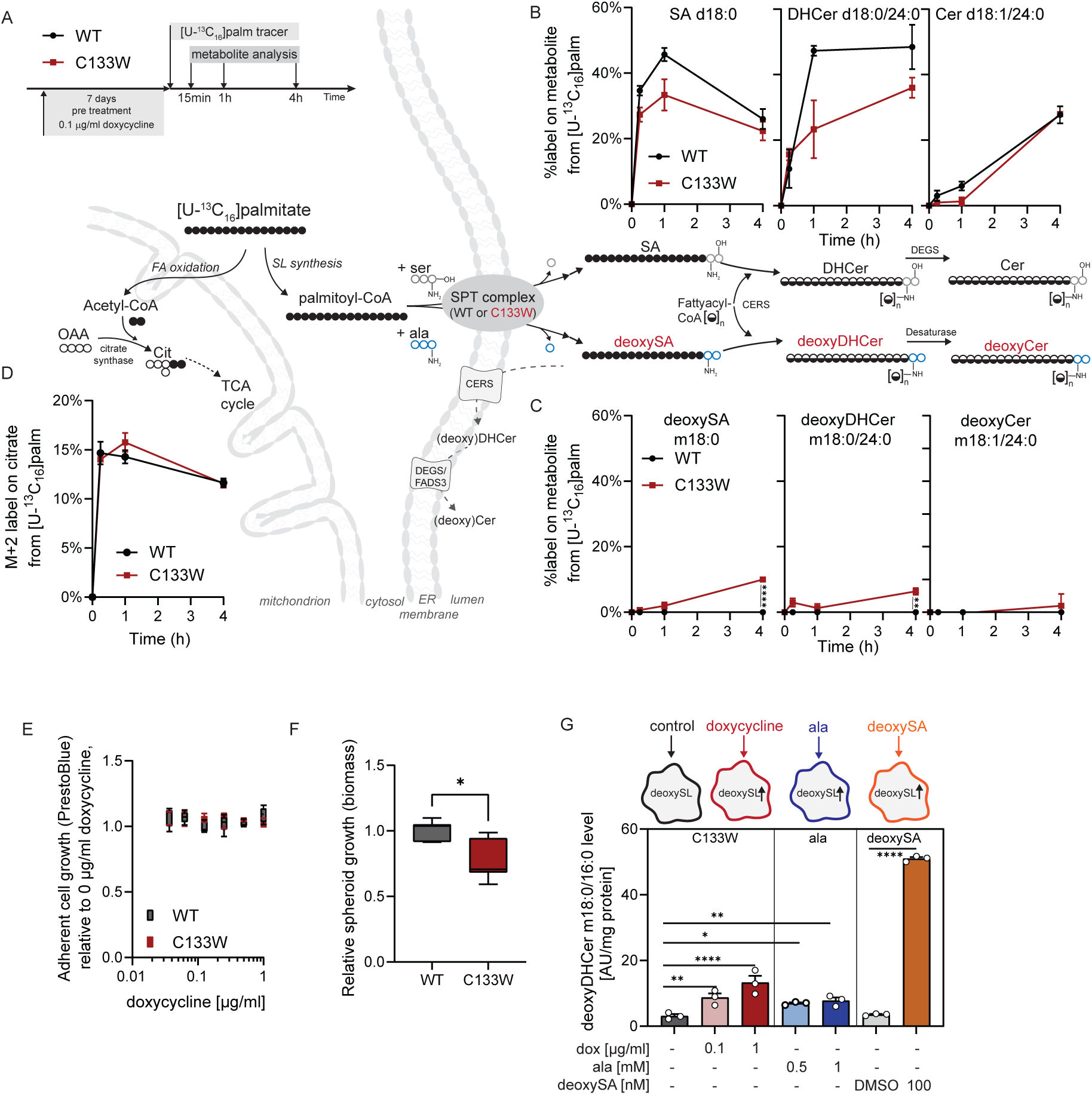
Impact of deoxySL synthesis on metabolic pathways. A. Schematic depicting trace into SL and deoxySL in engineered HCT116 cell lines cultured with 0.1 µg/ml doxycycline for 7 days prior tracing with [U-^13^C16]palmitate for 15min, 1h, and 4h. B. Label on canonical SLs from ^13^C palmitate tracer. C. Label on non-canonical SLs from ^13^C palmitate tracer. D. Label on citrate from ^13^C palmitate tracer. E. Adherent cell growth of *SPT* expressing HCT116 cell lines cultured for 3 days with increasing doxycycline concentrations. F. Spheroid growth in HCT116 *SPT* expressing cell lines with 0.1 µg/ml doxycycline. G. DeoxyDHCer m18:0/16:0 levels in HCT116 *SPTLC1^C133W^* expressing cells cultured for 3 days with alanine, doxycycline, or 100 nM deoxySA m18:0. Cells were grown in adherent (B-E, G) or spheroid cultures (F). Data are presented as means ± s.e.m. with three cellular replicates (B, C, D, G) or box (25^th^ to 75^th^ percentile with median line) and whiskers (min. to max. values) (E, F). Student’s *t*-test (B, D), nested *t*-test (F), or one-way ANOVA (G). Experiments in E, F, and G were repeated three times with similar results. Data in F depict average of 3 independent experiments with each 3 cellular replicates. \**P* < 0.05, ** *P* < 0.01, *** *P* < 0.001, **** *P* < 0.0001

We and others have also administered exogenous deoxySA to culture media for analysis of downstream effects (8,10,23), observing significant toxicity in some cell types. However, administration of free deoxySA significantly impacts its localization and thus local concentrations within cells compared to processing of newly synthesized deoxySLs on the ER membrane. In fact, we observed that deoxySA treatments, even at relatively moderate concentrations of 100 nM, resulted in a much greater accumulation of deoxyDHCer and deoxyCer species (∼20 fold) compared to alanine supplementation or *SPTLC1^C133W^* expression (2-5 fold) (Figs. 3G. S3K). These increases with deoxySA treatment were most pronounced in abundant deoxyDHCer species containing very-long chain fatty acids (Fig. S3L), highlighting that exogenous deoxysphingoid bases drive much higher deoxyDHCer biosynthetic flux compared to expression of the *SPTLC1^C133W^*. Notably, our adherent cell models achieved moderate, but more physiologically relevant, increases in deoxySL levels without showing overt cytotoxicity. Exogenous deoxySA administration may elicit distinct biological impacts due to aberrant localization of deoxySLs compared to that occurring via synthesis on the ER membrane. Indeed, ceramide synthases are expressed in the mitochondria and can drive cellular toxicity (36-38).

### Alanine and deoxySL synthesis influence colony formation in soft agar

Single-cell growth, migration, and invasive growth are hallmarks of transformed cells and correlate with anchorage-independent growth. As noted above, metabolically altering deoxySL synthesis consistently influenced “3D” anchorage-independent cell growth but not 2D adherent cell growth. To better understand the impact of deoxySL synthesis on proliferation in these contexts we next examined its impact on soft agar colony formation. Notably, we observed a significant growth defect in cells ectopically-expressing *SPTLC1^C133W^*compared to *SPTLC1^WT^* (Fig. 4A) with effects on colony size and number (Figs. S4A, B). We also rescued colony formation of *SPTLC1^C133W^*-expressing cells using 10 nM myriocin, which had little impact on colony formation in cells expressing *SPTLC1^WT^*(Fig. 4B). We also engineered the A549 non-small cell lung cancer cell line to express doxycycline-inducible *SPTLC1^WT^* or *SPTLC1^C133W^* (Fig. S4C). As in HCT116 cells, expression of *SPTLC1^C133W^*potently increased deoxySLs (Fig. S4D) and suppressed soft agar colony formation in these cells (Fig. S4E). On the other hand, myriocin treatment blocked growth suppression induced by *SPTLC1^C133W^* (Fig. S4E).

**Figure 4:**
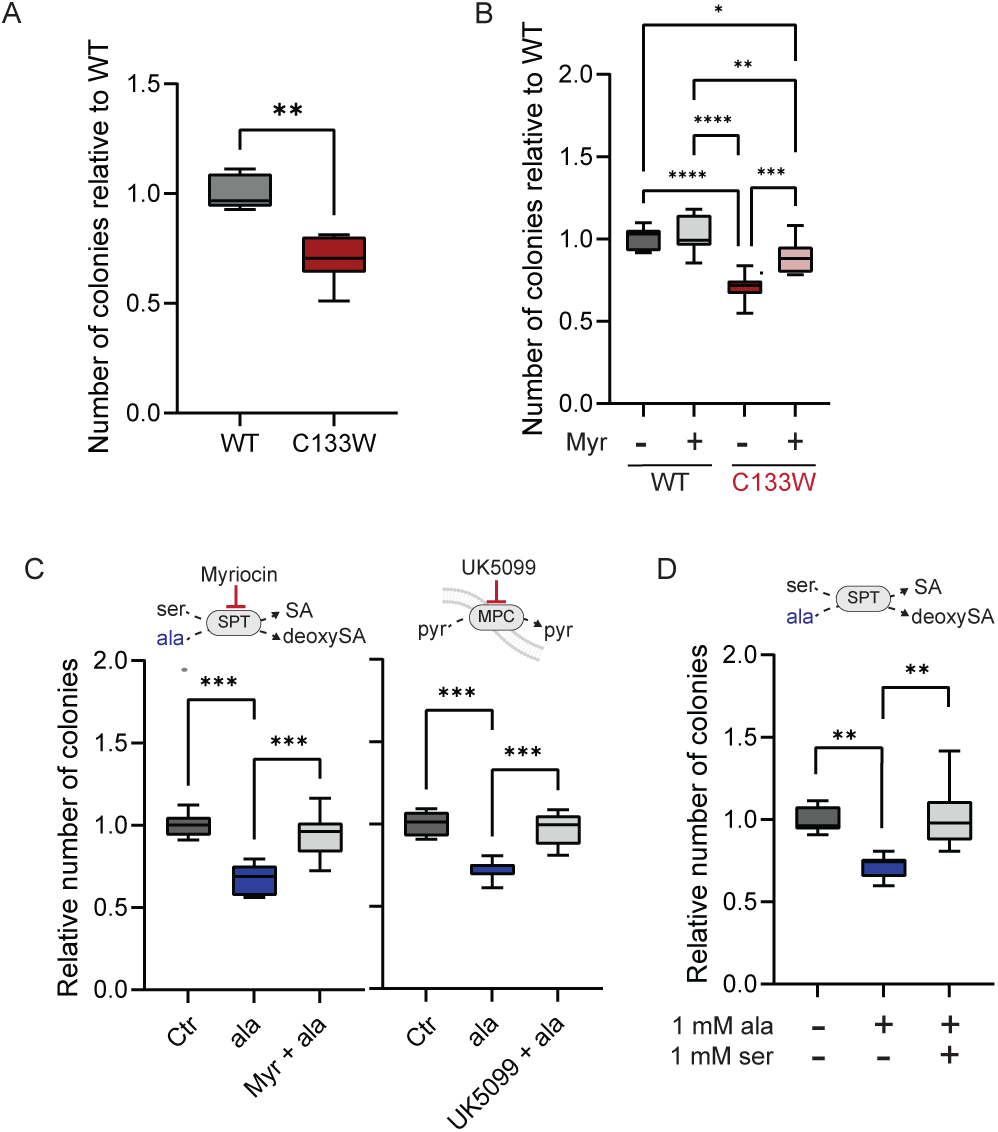
Alanine and deoxySL synthesis influence colony formation in soft agar. A. Colony formation in HCT116 wildtype (WT, grey) and mutant (C133W, red) *SPTLC1* expressing cells cultured for 7 days with 0.1 µg/ml doxycycline. B. Colony formation in *SPTLC1* HCT116 cells with 10nM myriocin cultured for 7 days with 0.1 µg/ml doxycycline. C. Colony formation in HCT116 cells with 1 mM alanine and 10 nM myriocin or 5 µM UK5099 cultured for 7 days with 0.1 µg/ml doxycycline. D. Colony formation in HCT116 cells with 1 mM alanine and 1mM serine cultured for 7 days. Cells were grown in adherent cultures and data are presented as box (25^th^ to 75^th^ percentile with median line) and whiskers (min. to max. values) obtained from three independent experiments with each three cellular replicates. Nested *t*-test (A) or nested one-way ANOVA (B-D) with \**P* < 0.05, ** *P* < 0.01, *** *P* < 0.001, \*\*\*\**P* < 0.0001.

To further support these results, we next examined whether buffering deoxySA synthesis induced by alanine supplementation rescues colony formation. Addition of alanine to culture media consistently reduced soft agar growth of HCT116 cells, but targeted inhibition of SPT or MPC, which both prevent deoxySL accumulation, mitigated the negative effects of this supplementation (Fig. 4C). Similar trends in soft agar colony formation were observed when modulating deoxySL synthesis in A549 cells with alanine and myriocin (Fig. S4F). Intriguingly, normalizing the serine/alanine ratio by co-addition of serine to the culture medium also enhanced soft agar growth, providing an orthogonal biochemical means of rescuing these effects (Fig. 4D). Collectively, these results suggest that metabolically induced deoxySL accumulation specifically impacts soft agar growth, or more generally, cellular processes important for anchorage-independent growth.

### DeoxySLs compromise plasma membrane endocytosis

The above studies demonstrate that deoxySL accumulation inhibits spheroid cell growth and soft agar colony formation. The temporal studies outlined in Figure 1F-G also highlighted the sensitivity of this process to cell seeding just after passaging, which induces membrane turnover facilitated by endocytosis. Soft agar growth also requires endocytic flux to catabolize matrix and establish multi-cellular colony formation. We therefore hypothesized that metabolic induction of deoxySL compromises plasma membrane endocytosis, which is mediated by GTPases such as RAB5. Since quantification and visualization of endocytosis and vesicular trafficking in 3D cultures are technically challenging, we determined the impact of deoxySL synthesis on endocytic events at the plasma membrane in adherent cell culture models. To directly examine whether this process was altered by deoxySL accumulation we quantified expression of the early endosomal marker RAB5, which regulates endosomal biogenesis and trafficking, in cells expressing *SPTLC1^WT^*or *SPTLC1^C133W^*. RAB5 positive endosomes were significantly decreased in *SPTLC1^C133W^*-expressing HCT116 cells compared to control conditions, indicating decreased endocytic activity (Fig. 5A). We also observed similar changes in A549 cells expressing *SPTLC1^C133W^*(Fig. 5B). Ectopic *SPTLC1^C133W^* expression induced reductions in RAB5 similar to that occurring with dynamin inhibition (Fig. 5A, B). Furthermore, we also observed decreased RAB5 positive endosomes in HCT116 cells induced to accumulate deoxySLs with alanine treatment, providing an additional link to defects in endocytosis (Fig. 5C). In fact, supplementation of both serine and alanine prevented this decrease in RAB5 positive endosomes from occurring (Fig. 5C).

**Figure 5:**
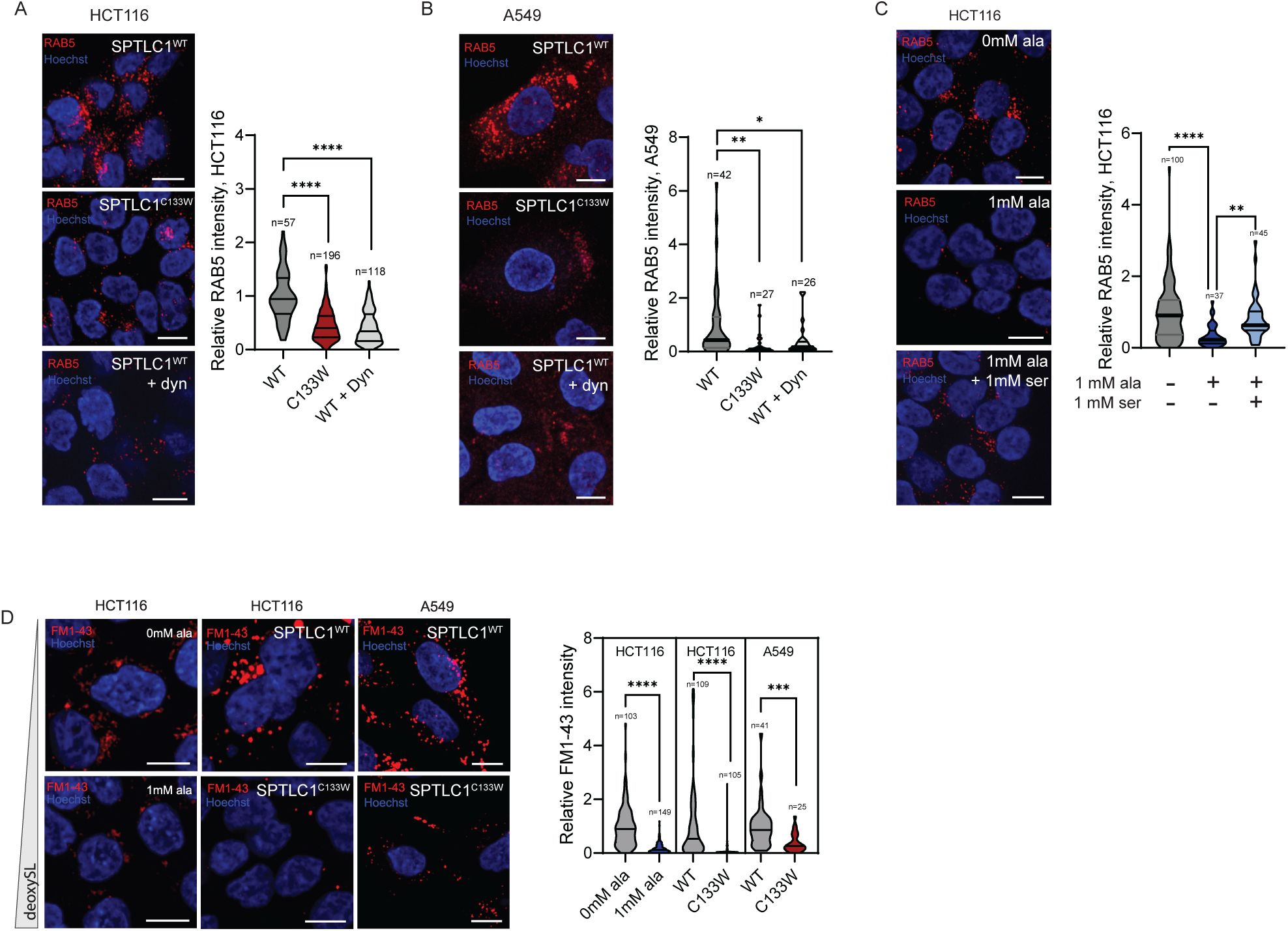
DeoxySLs compromise plasma membrane endocytosis. A. RAB5 immunostaining with *SPTLC1^WT^* and *SPTLC1^C133W^* expressing HCT116 cultured for 4 days with 0.1 µg/ml doxycycline. Dynasore (dyn) was given with 20 µM for 3h. B. RAB5 immunostaining with *SPTLC1^WT^* and *SPTLC1^C133W^* expressing A549 cultured for 4 days with 1 µg/ml doxycycline. Dynasore (dyn) was given with 20 µM for 3h. C. RAB5 immunostaining with HCT116 cells cultured with 1mM alanine or 1mM serine. D. FM1-43 staining in HCT116 and A549 cells cultured for 4 days with 0 or 1mM alanine or expressing wildtype (WT) or mutant (C133W) *SPTLC1*. Cells were grown in adherent cultures and violin blots depict relative RAB5 or FM1-43 intensity obtained from n = cell number. Data are representative of two independent experiments. White scale bar indicates 10 µm. One-way ANOVA (A, B, C) or student’s t-test (D) with \**P* < 0.05, ** *P* < 0.01, *** *P* < 0.001, **** *P* < 0.0001.

To further visualize alterations in membrane events, we imaged uptake of the styryl, membrane-impermeable lipid fluorescence probe FM1-43 and quantified internalized, FM1-43 stained vesicles as a measurement of endocytosis at the plasma membrane (39). Cells with high deoxySL synthesis, either induced by *SPTLC1*^C133W^ or alanine addition, had significantly decreased FM1-43 loading compared to control conditions (Fig. 5D), providing additional, functional evidence that deoxySL synthesis influences endocytic events at the plasma membrane.

## DISCUSSION

Here we investigated the effects of endogenous deoxySL accumulation in cancer cells, using alanine supplementation or ectopic expression of *SPTLC1^C133W^*. Our engineered cell systems allowed us to determine deoxySL-specific cellular effects independent of broader metabolic changes caused by serine and glycine deprivation. We observed that synthesis and accumulation of non-canonical deoxySLs specifically compromised anchorage-independent growth without influencing central carbon metabolism or respiration. Rather, these amino acid-driven metabolic changes influence plasma membrane endocytosis, highlighting an important functional role for SPT promiscuity in coordinating such processes (Fig. 6). While deoxySA synthesis has negligible effects on adherent cancer cell growth, we observed reduced proliferation in soft agar colony assays, an environment which places significant load on the plasma membrane. Therefore, in the context of dietary serine/glycine restriction, elevated deoxySLs may compromise tumor seeding and combine with additional downstream metabolic effects to reduce tumor progression (40). On the other hand, deoxySL synthesis also influences T-cell function (41), which also must be considered in the context of cancer therapy. Moving forward, these results clarify how deoxySL accumulation impacts cell metabolism and further highlight the impact of these species on membrane processes relevant for cancer cell, immune cell, and neuronal function.

**Figure 6:**
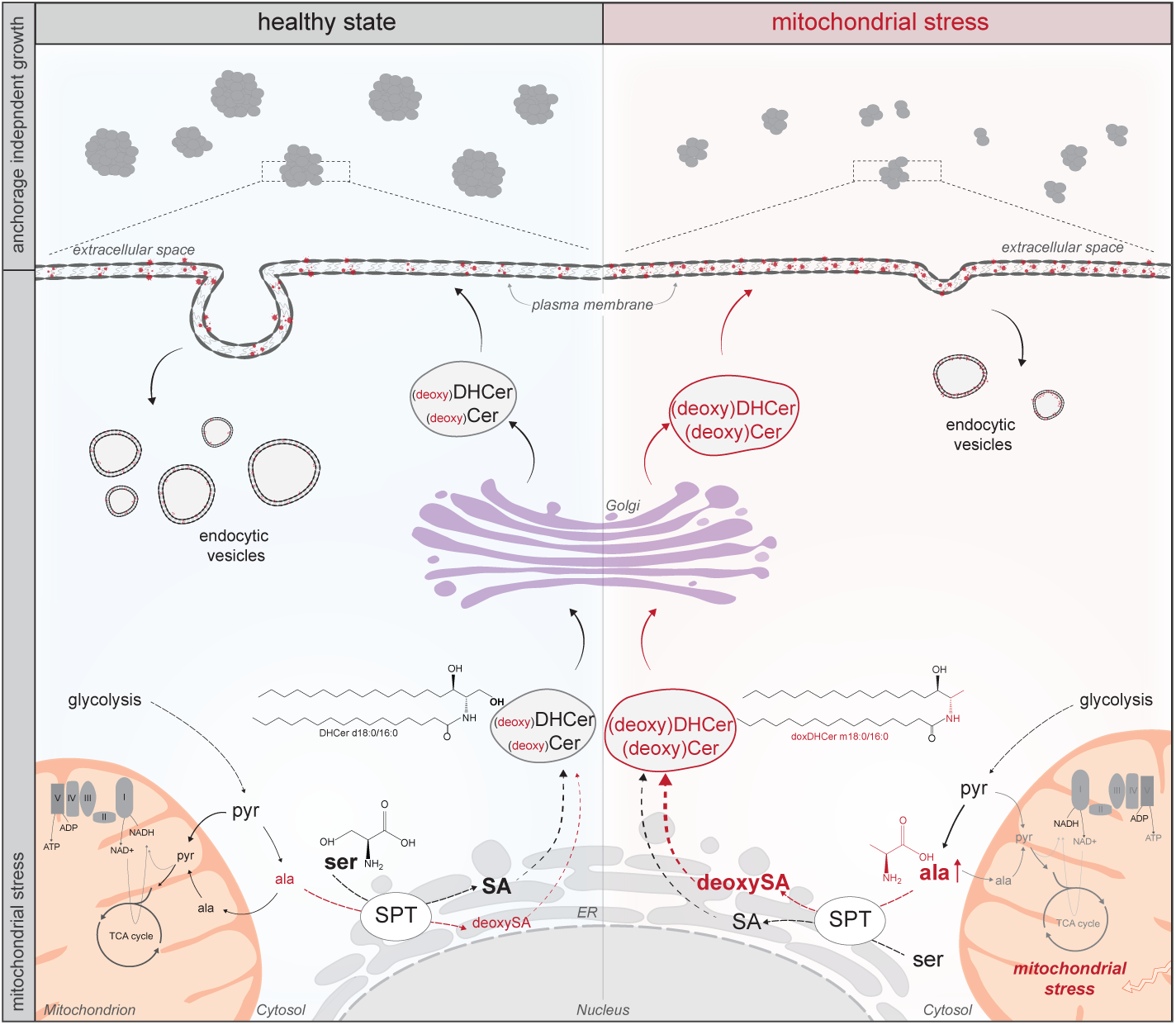
**Schematic metabolic concept** depicting metabolic impact of plasma membrane endocytosis through the lipidome compromising anchorage-independent cell growth. SPT – serine palmitoyltransferase.

We also observed differences in cells that synthesized deoxySLs endogenously versus those treated with exogenous deoxySA. Notably, SPT, CERS, dihydroceramide desaturase (DEGS), and fatty acid desaturase 3 (FADS3) are predominantly localized to the ER membrane, catalyzing the synthesis and processing of (deoxy)DHCer and (deoxy)Cer species that are further trafficked to membranes throughout the cell. Therefore, proteins and organelles exposed to deoxySA via exogenous treatments versus endogenous synthesis will differ, which could account for lack of any pronounced effect observed with *SPTLC1^C133W^*expression on intermediary metabolism.

Diverse phenotypes have been proposed for cells and tissues exposed to deoxySA, including mitochondrial dysfunction and apoptosis (8,20). However, HSAN1 patients acquire specific sensory defects with age and do not present with broad metabolic dysfunction (42), suggesting the effects of deoxySL accumulation are more subtle or specific to neurons. Rather, we observed specific effects of endogenously produced deoxySL on membrane processes that are critically important for anchorage-independent cancer cell growth and colony formation as well as neuronal function. As growth factor receptor-mediated signaling is intimately tied to plasma membrane trafficking, the impact on distinct signaling pathways (e.g. EGF or NGF) warrants further investigation (43,44). Furthermore, deoxySA regulation of sphingosine kinase could impact signaling through S1P receptors (22). Karsai et al. also recently observed that deoxySLs influence cell migration, which requires membrane dynamics for movement (20). Collectively these studies highlight key cellular phenotypes associated with deoxySL accumulation and membrane processes. One key question that remains is why the SPT complex retains this promiscuity and capacity for deoxySL synthesis. By integrating metabolic signals from amino acid metabolism into the ER and plasma membrane, SPT promiscuity could act as a regulatory node linking changes in amino acid or central carbon metabolism to lipid processing machinery.

## MATERIAL and METHODS

### Reagents

Media and sera were purchased from Life Technologies. Isotope tracers were purchased from Cambridge Isotopes. Sphingolipid standards were purchased from Avanti polar lipids. All other reagents were purchased from Sigma unless otherwise noted.

Cells were treated at a concentration of 5 µM UK5099 (Cat. #PZ0160, Millipore Sigma), 10 nM myriocin (Myr, Cat. #M1177, Millipore Sigma), 10 µM fumonisin B1 (FuB1, Cat. #F1147, Millipore Sigma), or 20 µM Fenofibrate (Cat. #PHR1246, Sigma) with stock solutions prepared in DMSO. DeoxySA m18:0 (Cat. #860493P, Avanti), and amino acids were supplemented to growth media with concentrations as indicated in the text.

### Cell culture

All cell lines were purchased from ATCC and tested negative for mycoplasma contamination: A549 (ATCC, CCL-185); HCT116 (ATCC, CCL-247); HEK293FT (ATTC, CRL-3216). Cells were cultured in Dulbecco’s Modified Eagle Media (DMEM, Cat. #11965-092, Gibco) containing 25 mM glucose, 4 mM glutamine, 100 U/ml penicillin, and 100 µg/ml streptomycin in a humidified cell culture incubator at 37 °C and 5% CO2. HCT116, A549, and HEK293T cells were cultured in growth medium containing 10% fetal bovine serum (FBS) (Cat. #16000-044, Gibco) and engineered *SPTLC1* expressing cell lines containing 10% tetracycline (TET)-free FBS (Cat. #FB15, Omega, Scientific. Inc.). Cells were detached with 0.05% trypsin-EDTA.

### Engineering SPTLC1 expressing cell lines

*SPTLC1* gene cDNA ORF nucleotide sequence (NM_006415) was purchased from GenScript. The ORF was subjected to point mutation to generate mutant *SPTLC1^C133W^* ORF and cloned into pCW57.1 vector (addgene Plasmid #41393). Briefly, the ORF and the plasmid were digested with Nhe1 and Age1, gel purified, and ligated. The engineered pCW57.1 plasmids containing *SPTLC1^WT^* or *SPTLC1^C133W^* were delivered into HCT116 and A549 cells using lentivirus delivery strategy.

Lentivirus were produced using HEK293T cells in high glucose DMEM supplemented with 10% FBS. One 10 cm cell culture dish of HEK293T cells at 60% confluency were transfected with 1.3 µg VSV.G/pMD2.G (addgene #12259), 5.3 µg lenti-gag/pol/pCMVR8.2 (addgene #12263), and 4 µg of the plasmid using 16 µL Lipofectamine 3000 diluted in 0.66 mL of OPTI-MEM (Life technology). Medium containing viral particles was harvested after 48 and 72 hours, filtered with 0.45 um filters, and further concentrated using Amicon Ultra15 centrifugal ultrafilters with a 100,000 NMWL cutoff (Cat. #UFC910024, Merck Millipore), and stored at -80°C until use.

Cells were infected in 6-well tissue culture plates with 6 µl concentrated virus particles in 0.5 ml growth medium containing 6 µg/ml polybrene for 4h before addition of 2 ml virus-free growth medium containing polybrene. After 24h, medium was changed to growth medium containing 2 µg/ml puromycin. After puromycin selection, immunoblotting and RT-PCR analyses were carried out to confirm the expression of SPTLC1 in response to doxycycline as indicated in the text. For metabolic and functional experiments, HCT116 cells were induced with 0.1 µg/ml doxycycline and A549 cells with 1 µg/ml doxycycline.

### Adherent cell growth assay using PrestoBlue

Cells were cultured in 96 well plates for 3 days with treatments as indicated in each figure. Cell viability was determined using PrestoBlue Cell viability Reagent (Invitrogen) per manufacturer’s instructions.

### Spheroid growth assay

For spheroid assays, 20,000 singularized cells were plated in 2 ml growth medium on 6-well ultra-low attachment plates (Cat. #3471, Corning) with treatments as indicated in the text (1 mM alanine, 5 µM UK5099, 10 nM myriocin, 20 µM fumonisin B1). HCT116 expressing cell lines were induced with 0.1 µg/ml doxycycline for 2 days prior seeding. Medium was changed on day 3 and 5. Growth was determined after 3-8 days as indicated in the text. Spheroids were collected in 2 ml sample tubes, centrifuged at 100 *g* for 5 min and the pellet was washed once with 2 ml 0.9% (w/v) NaCl. The spheroids were lysed with 100 µl RIPA buffer (Cat. #BP-115-5x, Boston Bioproducts) and homogenized with a ball mill (Retsch Mixer Mill MM400) at 30Hz for 3 min. The cell suspension was centrifuged at 20,000 *g* for 10 min at 4 °C and 20 µl of the supernatant was used for protein quantification using BCA protein assay kit (Cat. #G1002, Lamda Biotech. Inc) as per manufacturer’s instruction.

### Soft agar colony formation assay

Soft agar colony formation assay was conducted in 6-well tissue culture plates. A 3% low gelling temperature agarose (Cat. #A4018, Millipore Sigma) stock solution was prepared in water, autoclaved, and kept at 55 °C in a water bath until use. 2 ml of the lower, hard layer containing of 0.6% in growth medium was added per well and allowed to solidify for 20 min at room temperature. 3000 cells were mixed with 1 ml of 0.3% agarose in growth medium supplemented with treatments of choice (DMSO, 5 µM UK5099, 10 nM myr, 1 mM serine, 0.5 mM or 1 mM alanine) as indicated in figure legends and overlayed on the lower, hard layer. The plates were placed at 4°C for 5 min to allow for a fast solidification improving uniform cell distribution in the soft agarose layer. The plates were then transferred into a humidified cell culture incubator at 37 °C with 5% CO2 to allow colony formation. HCT116 colonies were counted on day 7 using a brightfield microscope. Pictures of soft agar colonies were taken with a Leica DMi8 Microsystem with 5x, 10x, or 20x objective as indicated in each figure legend. Colonies were stained by addition of 0.3 ml of 0.1 % crystal violet solution per well and incubated for 15 min at room temperature. The wells were washed five times with 1 ml water and incubated for 5 min each at room temperature. Pictures were taken with the BioRad ChemiDoc XRS+ imaging station.

### Protein extraction and immunoblotting

Cells were plated on 6-well plates containing in growth medium containing 10% TET-free FBS and doxycycline for 4 days as indicated in the figure legend. Medium was changed on day 2. Cells were lysed in ice-cold RIPA buffer (Cat. #BP-115-5x, Boston Bioproducts) supplemented with 1x HALT protease inhibitor cocktail (Cat. #78430, Thermo Fisher Scientific), vortexed for 20 min at 4°C and centrifuged at 20,000 *g* for 10 min at 4 °C. Supernatant was used to quantify protein using BCA protein assay kit (Cat. #G1002, Lamda Biotech. Inc) and pre-diluted Protein Assay Standards (Cat. #23208, Thermo Scientific) as per manufacturer’s instruction. Samples were incubated for 5 min at 95 °C in 4x laemmli buffer. 40 μg of total protein was separated on a 4–20% SDS-PAGE gel (mini-PROTEAN TGX gels, Bio-Rad) along with PageRuler Prestained protein ladder (Cat. #26619, Thermo Scientific) and proteins were transferred onto a PVDF membrane (Millipore, Cat. No. ISEQ00010). The membrane was blocked with 5% non-fat milk in tris buffered saline with 0.1% Tween 20 (TBS-T) for 1 h and immunoblotted with primary antibody at 4 °C overnight diluted in 5% non-fat milk. anti-SPTLC1 (1:1000 dilution, Cat. #15376-1-AP, Proteintech), anti-ß-actin (1:1000 dilution, Cat. #3700, Cell Signaling Technology). The immunoblots were then incubated with secondary antibody for 1 h at room temperature (1:1000 dilution, anti-rabbit Cat. #7074, lot 28, or anti-mouse HRP-conjugate Cat. #7076S, lot 32, Cell Signaling Technology). Specific signal was detected using SuperSignal West Pico Chemiluminescent Substrate (Cat. # 1705061, BioRad) and imaged with a BioRad ChemiDoc XRS+ imaging station. Band signal from Western Blots was determined using Image Lab Software from BioRad and normalized to signal of ß-actin. Antibodies for western blotting were validated for human reactivity by the manufacturer and used per their instructions. Uncropped raw blots are provided in Extended Data 1.

### Respirometry

Respiration was measured in adherent monolayers of cells using an Agilent Seahorse XFe96 Analyzer with a minimum of 5 cellular replicates per condition. Engineered *SPTLC1^C133W^* expression cell line were cultured in the presence of doxycycline four days prior and two days after seeding on 96well plates in the presence of doxycycline. Cells were assayed in DMEM medium (Cat. #5030, Sigma Aldrich) supplemented with 8 mM glucose, 3 mM glutamine, 3 mM pyruvate, and 2 mM HEPES. Cells were washed twice with 100 µl assay medium and cultured in 150 µL assay medium for 1h before measurement. Respiration was measured under basal conditions as well as after injection of 2 μM oligomycin (Oligo) (Port A), sequential addition of 200 nM FCCP (Port C, D), and addition of 0.5 μM rotenone and 1 μM antimycin (Ant/Rot) (Port D). Oxygen consumption rates were normalized to protein content using BCA protein assay kit (Cat. #G1002, Lamda Biotech. Inc).

### Isotopic tracing and analysis

Cells were cultured in medium containing stable isotope tracers of choice as indicated in the text. Tracers were purchased from Cambridge Isotopes Inc. or Sigma: [U-^13^C_6_]glucose (CLM-1396-25), [U-^13^C16]palmitate (CLM-409), and [2,3-^13^C2]alanine (Cat. #604682). For alanine tracing studies, HCT116 cells were cultured in growth medium supplemented with 1 mM ^13^C alanine for 24h in the presence of DMSO or 5 µM UK5099. For glucose isotopic labeling experiments, HCT116 *SPTLC1* expressing cells were induced with 0.1 µg/ml doxycycline for 5 days before starting tracing for 24h. Tracing was performed with DMEM (Cat. #5030, Sigma Aldrich) medium where ^12^C glucose replaced with 25 mM [U-^13^C_6_]glucose and 10% TET-free FBS was supplemented. Labeling on metabolites from ^13^C alanine and ^13^C glucose was quantified using GC-MS technology. Mass isotopomer distributions and total metabolite abundances were computed by integrating mass fragments using a MATLAB based algorithm with corrections for natural isotope abundances as described previously (45,46).

For tracing studies with [U-^13^C16]palmitate, HCT116 cells were cultured in growth medium in the presence of 0.1 µg/ml doxycycline for 7 days before tracer start. Growth medium was replaced to DMEM medium containing 1% (v/v) delipidated FBS 24h prior tracer start and medium exchange again 1h prior tracer trace. [U-^13^C16]palmitate was noncovalently bound to fatty acid-free BSA and added to culture medium at 5% of the final volume (50 μM final concentration). Media was prewarmed to 37°C in a cell incubator with 5% CO2 and cells were traced for 15 min, 1 h, and 4 h. Metabolites were extracted, and labeling was quantified on a Q-Exactive LC-MS system. Metabolites were annotated and labeling was corrected for natural isotope abundance using Maven software v2011.6.17 (47). Due to [U-^13^C16]palmitate tracer purity of 98% (Cat#. #CLM-409, Cambridge Isotope Laboratories, Inc), we also observed a low fraction of labeled M+15 as well as M+31 on deoxyDHCer m18:0/16:0. In Figures S3A and S3B, M+16 depicts the combined label of M+15 and M+16, while M+32 depicts the combined label of M+31 and M+32.

### Gas chromatograph - Mass spectrometry (GC-MS) and sample preparation

Metabolites were extracted, analyzed, and quantified, as previously described in detail (46). Briefly, cells were washed with saline solution and quenched with 0.25 ml -20 °C methanol. After adding 0.1 ml 4°C cold water, cells were collected in tubes containing 0.25 ml -20 °C chloroform. The extracts were vortexed for 10 min at 4 °C and centrifuged at 16,000 × *g* for 5 min at 4°C. The upper aqueous phase was evaporated under vacuum at 4 °C. Derivatization for polar metabolites was performed using a Gerstel MPS with 15 μl of 2% (w/v) methoxyamine hydrochloride (Thermo Scientific) in pyridine (incubated for 60 min at 45°C) and 15 μl N-tertbutyldimethylsilyl-N-methyltrifluoroacetamide (MTBSTFA) with 1% tert-butyldimethylchlorosilane (Regis Technologies) (incubated further for 30 min at 45 °C). Derivatives were analyzed by GC-MS using a DB-35MSUI column (30 m x 0.25 i.d. x 0.25 μm) installed in an Agilent 7890B gas chromatograph (GC) interfaced with an Agilent 5977A mass spectrometer (MS) operating under electron impact ionization at 70 eV. The MS source was held at 230°C and the quadrupole at 150 °C and helium was used as carrier gas. The GC oven was held at 100°C for 2 min, increased to 300 °C at 10 °C/min and held for 4 min, and held at 325°C for 3 min.

### FM^TM^ 1-43 dye uptake assay, RAB5 immunostaining, and fluorescence microscopy

Cells were plated on 12-well plates containing cover glass coverslips for 4 days in the presence of doxycycline or small molecule treatments as indicated in the text. Medium was changed on day 2. Media was supplemented with 20 µM dynasore (Cat. #D7693, Sigma Millipore) 3 h. To visualize RAB5 positive endosomes, cells were fixed with ice-cold 4% paraformaldehyde solution (Cat. #J19943-K2, Thermo Scientific), washed three times with 0.1% Tween-20 in PBS (PBST) before incubation with 0.2% Triton-X100 for 10 min. Cells were then washed 3 times with PBST (each 5 min), incubated with 8% BSA in PBST (Cat. #700-100P, Gemini Bio-Products) for 20 min, and washed three times with PBST (each 5 min). Cells were then incubated with primary anti-RAB5 antibody at 4 °C overnight diluted in 8% BSA in PBST (1:500 dilution, Cat. #3547, lot 7, Cell Signaling Technology). Secondary anti-rabbit Alexa Fluor 568 antibody (1:1000 dilution, Cat. #A-11011, Life Technologies) and Hoechst (1:1000 dilution, Cat. #4082, Cell Signaling Technology) diluted in 8% BSA PBST were applied for 2 h at room temperature followed by an incubation of HCS Cell Mask Deep Red Stain (Cat. #H32721, Invitrogen) for 30 min at room temperature. Cells were then washed three times with PBST (5 min each). Antibodies were validated for human reactivity by the manufacturer and used per their instructions. Coverslips were mounted with Fluoromount-G (Cat. #0100-01, Southern Biotech) on microscopy plates (Cat. #12-544-7, Thermo Fisher Scientific), and sealed with nail polish (Cat. #88-128, Genesee Scientific Corp.).

For plasma membrane staining, cells were incubated with 5 µg/ml FM^TM^ 1-43FX (Cat. #F35355, Invitrogen) for 10 min at 37°C and fixed with ice-cold 4% paraformaldehyde solution as per manufacturer’s instruction. Nuclei were visualized with Hoechst as described above.

Cells were imaged using a 60x Plan Apo 1.4 NA objective on a Nikon Ti2-E microscope with a Yokogawa X1 spinning disk confocal system, MLC400B 4-line (405nm, 488nm, 561nm, and 647nm) dual-fiber laser combiner (Agilent), Prime 95B back-thinned sCMOS camera (Teledyne Photometrics), piezo Z-stage (Mad City Labs) and running NIS Elements software. To separate the emission of individual flours, band-pass emission filters were used for each channel (450/50, 525/36, 605/52, and 705/72). Background fluorescence intensity was corrected for each image. One representative image is depicted per experimental condition. RAB5 was quantified using ImageJ 1.53k with n cell number as depicted in each figure.

### Liquid chromatograph - Mass spectrometry (LC-MS), sample preparation, and sphingolipid analysis

For targeted sphingolipid analysis, cells were washed with 0.9% (w/v) NaCl and extracted with 0.25 mL of −20°C methanol, 0.25 mL -20°C chloroform, and 0.1 mL of water spiked with deuterated internal standards (20 picomoles of D7-sphinganine (Avanti, Croda International Plc, 860658), 2 picomoles of D3-deoxysphinganine (Avanti, Croda International Plc, 860474), 200 picomoles of C15 ceramide-d7 (d18:1-d7/15:0) (Avanti, Croda International Plc, 860681), 100 picomoles of C13-dihydroceramide-d7 (d18:0-d7/13:0) (Avanti, Croda International Plc, 330726)). The tubes were vortexed for 5 min, centrifuged at 20,000 x *g* at 4°C for 5 min, and the lower organic phase was collected. The remaining polar phase was re-extracted with 2 μL formic acid and 0.25 mL of -20 °C chloroform. The organic phases were combined, dried under air, resuspended in 80 μL Buffer B (0.2% formic acid and 1 mM ammonium formate in methanol), sonicated for 10 min, and centrifuged for 10 min at 20,000 x *g* at 4 °C. Ceramide species in the supernatant were quantified using LC-MS (Agilent 6460 QQQ, MassHunter LC/MS Acquisition (v.B.08.02)) equipped with a C8 column (Spectra 3 μm C8SR 150 × 3 mm inner diameter, Peeke Scientific) as previously described (Bielawski et al., 2009) with mobile phase A (2 mM ammonium formate and 0.2% formic acid in HPLC grade water) and mobile phase B (0.2% formic acid and 1 mM ammonium formate in methanol). The gradient elution program consisted of the following profile: 0 min, 82% B; 3 min, 82% B; 4 min, 90% B, 18 min, 99% B; 25 min, 99%, 27 min, 82% B, 30 min, 82% B. Column re-equilibration followed each sample and lasted 10 min. The capillary voltage was set to 3.5 kV, the drying gas temperature was 350 °C, the drying gas flow rate was 10 L/min and the nebulizer pressure was 60 psi. Ceramide species were analyzed by selective reaction monitoring of the transition from precursor to product ions at associated optimized collision energies and fragmentor voltages are provided elsewhere (10). SL and deoxySL species were then quantified from spiked internal standards and normalized to protein content.

To quantify labeling on SL and deoxySL species from [U-^13^C16]palmitate a Q Exactive orbitrap mass spectrometer with a Vanquish Flex Binary UHPLC system (Thermo Scientific) was used with a Kinetex 2.6 µM C8 100 Å 150 x 3 mm LC column (Phenomenex) at 40°C. 5 μL of sample was injected. Chromatography was performed using a gradient of 2 mM ammonium formate and 0.2 % formic acid (mobile phase A) and 1 mM ammonium formate and 0.2 % formic acid in methanol (mobile phase B), at a flow rate of 0.5 mL/min. The LC gradient held at 82% B for 0-3 min, then ran from 82%-90% B in 3-4 min, then 90-99% in 4-18 min, held at 99% B for 7 min, then reduced from 99%-82% from 25-27 min, then held at 82% for a further 13 mins. Lipids were analyzed in positive mode using spray voltage 3 kV. Sweep gas flow was 5 arbitrary units, auxiliary gas flow 7 arbitrary units and sheath gas flow 50 arbitrary units, with a capillary temperature of 300°C. Full MS (scan range 150-2000 m/z) was used at 70 000 resolution with 1e^6^ automatic gain control and a maximum injection time of 200 ms. Data dependent MS2 (Top 6) mode at 17 500 resolution with automatic gain control set at 1e^5^ with a maximum injection time of 50 ms was used for peak identification, combined with known standards where possible. Abundances of SL and deoxySL species were normalized to the spiked internal standards and protein content.

### Statistics

Data visualization and statistical analysis were performed using GraphPad Prism (v9.1.0) and Adobe Illustrator CS6 (v.16.0.0). The type and number of replicates and the statistical test used are described in each figure legends. Data are presented as means ± s.e.m. or box (25^th^ to 75^th^ percentile with median line) and whiskers (min. to max. values). Tissue culture was conducted in 12-well tissue culture plates for metabolic studies, 6-well tissue culture plates for soft agar colony formation assay, and 96-well culture plates for Seahorse analysis and growth responses (PrestoBlue assay). Each culture well was considered as a cellular replicate for a given preparation. One representative figure per condition is depicted for confocal microscopy analysis. *P* values were calculated using a two-sided Student’s *t*-test, one-way ANOVA or two-way ANOVA, and nested-analysis was performed as indicated, with \**P* < 0.05; \*\**P* < 0.01; \*\*\**P* < 0.001, and \*\*\*\**P* < 0.0001 as indicated in each figure legend.

## Data availability

The datasets generated and analyzed during the current study are available in the article and are available from the corresponding author upon reasonable request. The SPTLC1^WT^ and SPTLC1^C133W^ mammalian expression constructs are available at Addgene (addgene #181920 and #181921). Data obtained from high-resolution mass spectrometry are available at the Metabolomics Workbench.

## Acknowledgments

We thank the Nikon Imaging Center at UC San Diego, where we collected imaging data.

## Authors’ contribution

C.M.M. and T.C. designed the study. T.C. performed *in vitro* cell studies and targeted metabolomics. T.C., T.M., and J.G. engineered *SPTLC1* expression cells. T.M. performed time-dependent spheroid experiments. T.C. and G.H.MG. performed high-resolution mass spectrometry. T.C. and R.S.K. performed fluorescence imaging experiments. T.C. and S.V.K. performed soft agar growth assays. T.C. analyzed data and prepared figures. C.M.M. and T.C. wrote the manuscript with input from all authors.

## Abbreviations

[U-^13^C16]palmitat: Uniformly labeled palmitate containing 16 isotopic labeled carbons
Cer: Ceramides
CERS: Ceramide synthases
DEGS: Dihydroceramide desaturases
deoxyCer: Deoxy-ceramides
deoxyDHCer: Deoxy-dihydroceramides
deoxySA: 1-deoxy-sphinganine
deoxySL: 1-deoxysphingolipid
DHCer: Dihydroceramides
ER: Endoplasmic reticulum
FADS3: Fatty acid desaturase 3
FuB1: Fumonisin B1
HSAN1: Hereditary sensory neuropathy type I
Myr: Myriocin
SL: Sphingolipid
SPT: Serine palmitoyltransferase
*SPTLC1^C133W^*: Mutant C133W serine palmitoyltransferase subunit LC1
*SPTLC1^WT^*: Wildtype serine palmitoyltransferase subunit LC1

---

**Supplementary Figure 1:**
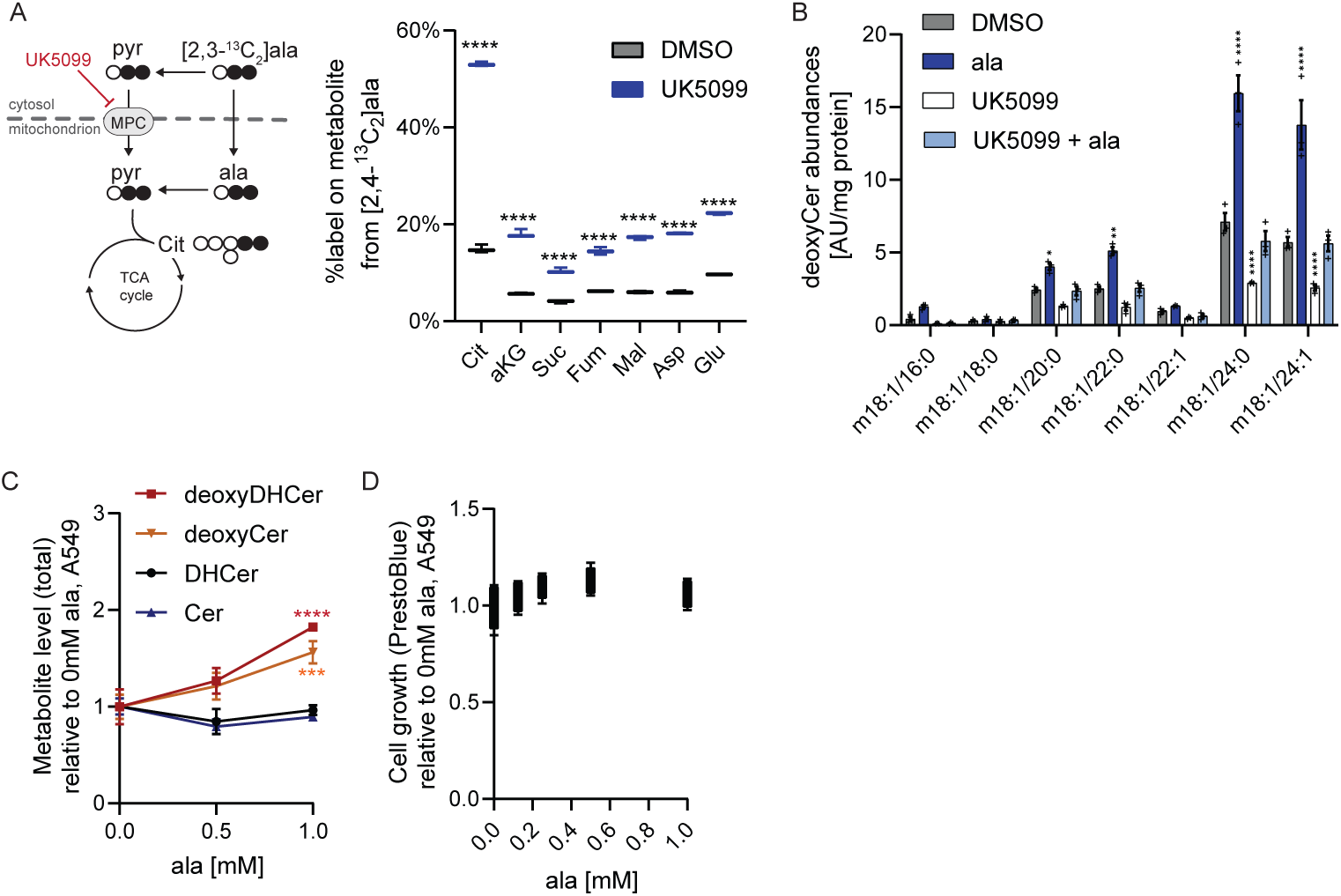
Alanine promotes deoxySL synthesis. A. Schematic depicting [2,3-^13^C]alanine trace into TCA cycle metabolism. Open circles represent ^12^C, closed circles ^13^C atoms. Percent label on TCA cycle intermediates in HCT116 cells cultured with UK5099 and ^13^C alanine for 24h. B. Levels of deoxyCer species in HCT116 cells cultured for 3 days with 1mM alanine and 5 µM UK5099. C. Total deoxyDHCer, deoxyCer, DHCer, and Cer levels in A549 cells cultured with increasing alanine for 3 days relative to 0 mM alanine. D. Adherent cell growth of A549 cells cultured with increasing alanine concentrations for 3 days. Data are presented as means ± s.e.m. (B, C) or box (25^th^ to 75^th^ percentile with median line) and whiskers (min. to max. values) (A, D) with three (A, B, C) or five (D) cellular replicates. Student’s t-test (A) or two-way ANOVA (B, C). Significance in A is compared to 0mM alanine. Significance in B is compared to DMSO control condition. Results are depicted from one representative experiment and each experiment was repeated independently three times with similar results. * *P* < 0.05, ** P < 0.01, *** *P* < 0.001, **** *P* < 0.0001.

**Supplementary Figure 2:**
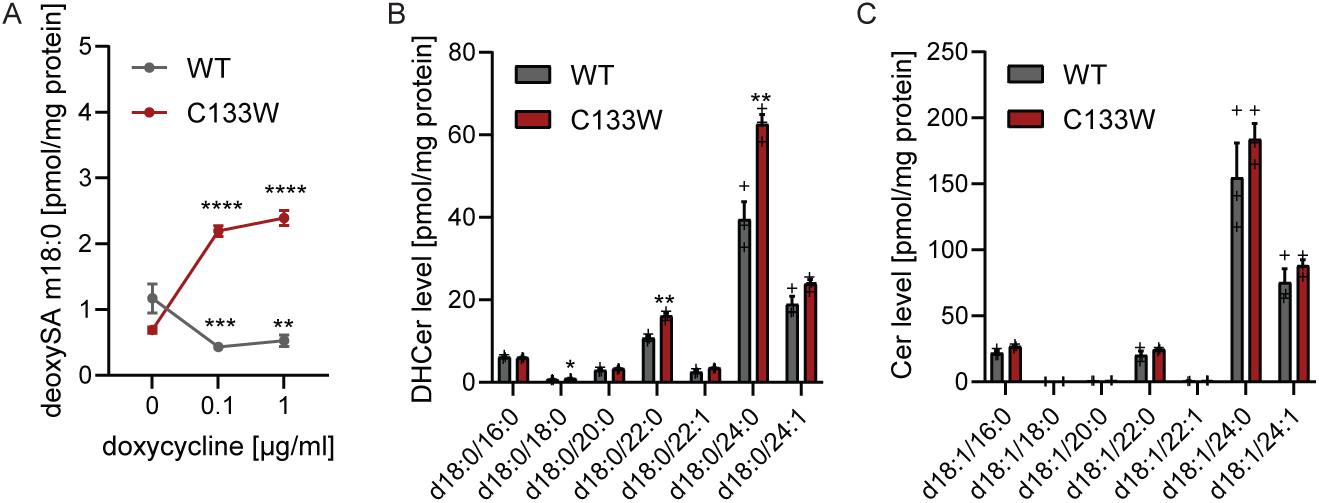
Mutant *SPTLC1* drives non-canonical deoxySL biosynthesis. A. DeoxySA levels in SPT expressing HCT116 cells. B. Levels of DHCer species in HCT116 cell lines. C. Levels of Cer species in HCT116 cell lines. Cells were cultured for 5 days with 0.1 µg/ml doxycycline. Data are presented as means ± s.e.m. with three cellular replicates. Results are depicted from one representative experiment and each experiment was repeated independently three times with similar results. Two-way ANOVA (A) relative to 0µg/ml doxycycline or Students *t*-test (B,C) with \**P* < 0.05, ** *P* < 0.01, *** *P* < 0.001, **** *P* < 0.0001.

**Supplementary Figure 3:**
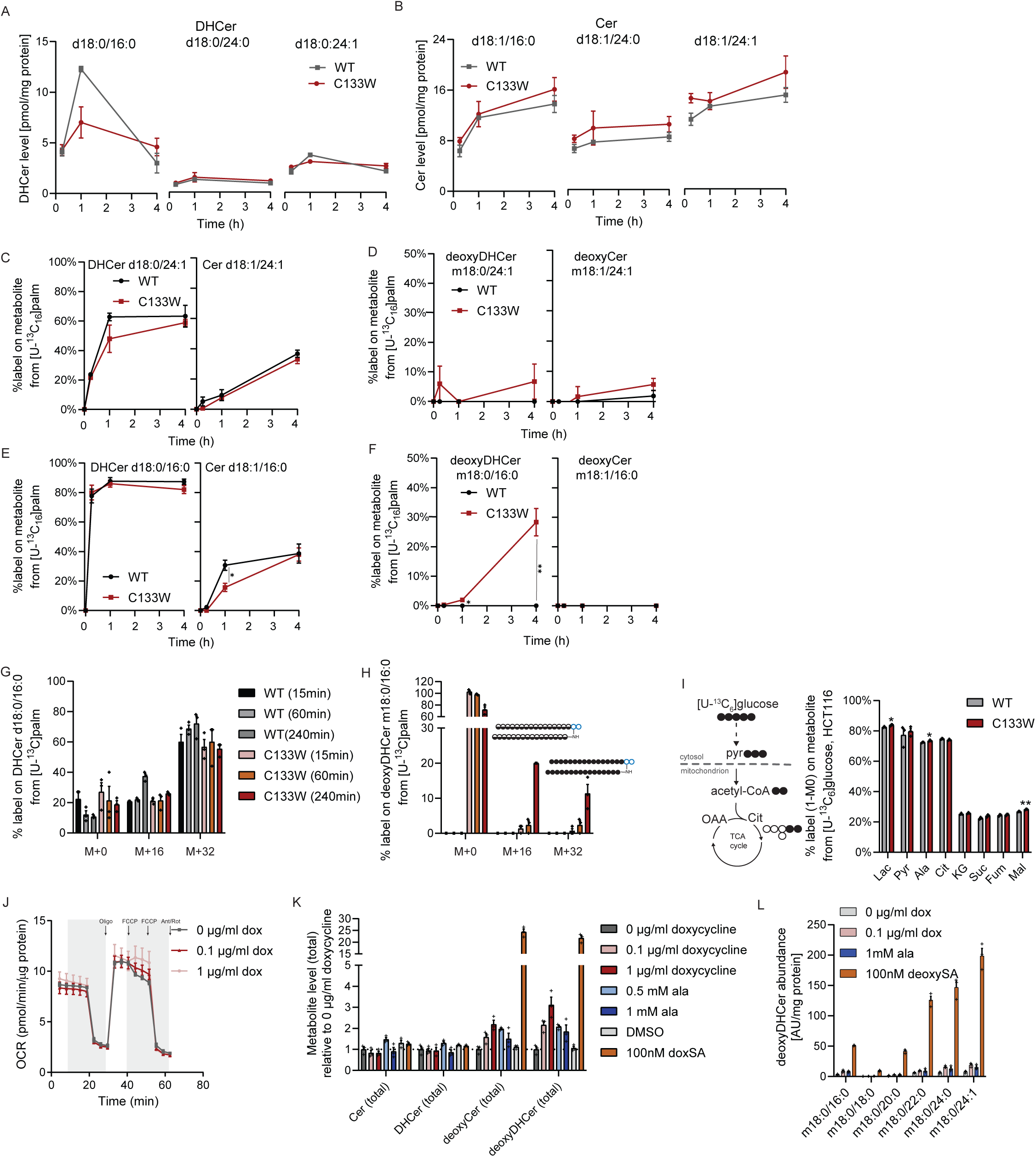
Impact of deoxySL synthesis on metabolic pathways. A. Abundances of DHCer species in HCT116 cells. B. Abundances of Cer species in HCT116 cells. C. Label on canonical 24:1 species from ^13^C palmitate tracer in HCT116 cells. D. Label on non-canonical 24:1 species from ^13^C palmitate tracer in HCT116 cells. E. Label on canonical 16:0 species from ^13^C palmitate tracer in HCT116 cells. F. Label on non-canonical 16:0 species from ^13^C palmitate tracer in HCT116 cells. G. Percent label on deoxyDHCer m18:0/16:0 from ^13^C palmitate tracer in HCT116 *SPTLC1^C133W^* expressing cells. H. Percent label on DHCer d18:0/16:0 from ^13^C palmitate tracer in HCT116 *SPTLC1^C133W^* expressing cells. I. Atom transition map depicting [U-^13^C_6_]glucose oxidation with ^12^C (open circles) and ^13^C (closed circles) carbons into TCA cycle. Labeling on metabolites in SPT expressing HCT116 cells cultured for 4d with 0.1 µg/ml doxycycline followed by 48h trace with [U-^13^C_6_]glucose. J. Oxygen consumption (OCR) in HCT116 SPTLC1^C133W^ cells cultured for 6 days with increasing doxycycline concentrations. K. DeoxyDHCer species in HCT116 *SPTLC1^C133W^* cells cultured for 3 days with doxycycline, alanine or deoxySA m18:0. L. Levels of total sphingolipid species in HCT116 *SPTLC1^C133W^* cells cultured for 3 days with doxycycline, alanine, and deoxySA m18:0. Data are presented as means ± s.e.m. with three (A-I, K, L) or five (J) cellular replicates. Results are depicted from one representative experiment and each experiment was repeated independently two or more times with similar results. Results in A -H were obtained from one independent experiment. Students *t*-test with \**P* < 0.05, ** *P* < 0.01, *** *P* < 0.001.

**Supplementary Figure 4:**
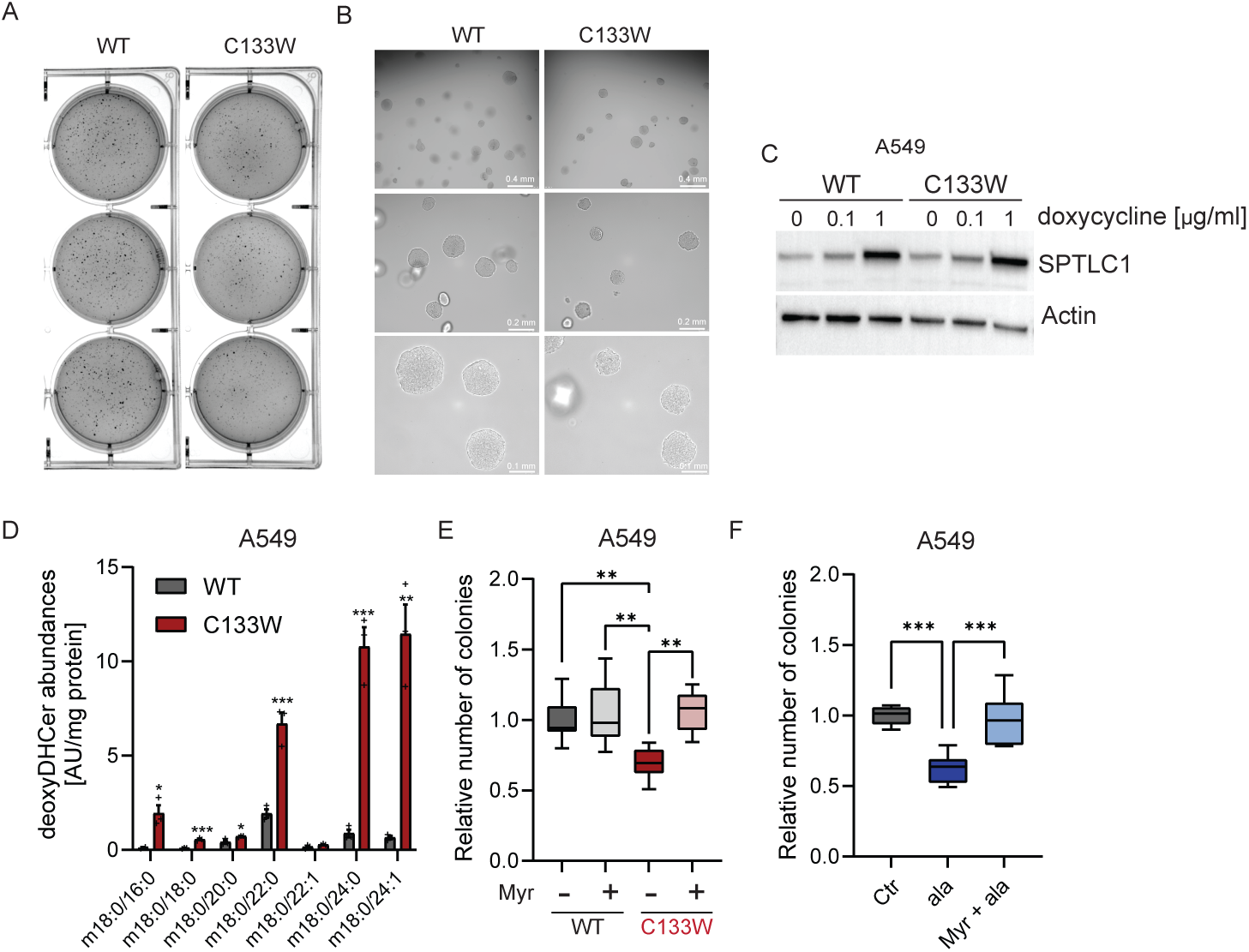
Impact of deoxySL synthesis on colony formation. A. Representative soft agar colony pictures of HCT116 cells expressing wildtype (WT) or mutant (C133W) *SPTLC1* grown in soft agar for 7 days with 0.1 µg/ml doxycycline stained with crystal violet staining. B. Representative soft agar colony pictures of HCT116 cells expressing wildtype (WT) or mutant (C133W) *SPTLC1* grown with 0.1 µg/ml doxycycline C. Immunoblotting of SPTLC1 and actin in *SPTLC1* expressing A549 cells with doxycycline for 4 days. D. Levels of deoxyDHCer species in engineered A549 cells cultured for 4 days with 1 µg/ml doxycycline. E. Colony formation in engineered A549 cells cultured for 14 days with 10 nM myriocin (Myr) and 1 µg/ml doxycycline. F. Colony formation in A549 cells cultured for 14 days with 1 mM alanine and 10 nM myriocin. Data are presented as means ± s.e.m. with three cellular replicates (D). Data in E and F are presented as box (25^th^ to 75^th^ percentile with median line) and whiskers (min. to max. values) obtained from three independent experiments with each three cellular replicates. Student’s t-test (D) or nested one-way ANOVA (E, F). \**P* < 0.05, ** *P* < 0.01, *** *P* < 0.001, **** *P* < 0.0001

**Extended data 1:**
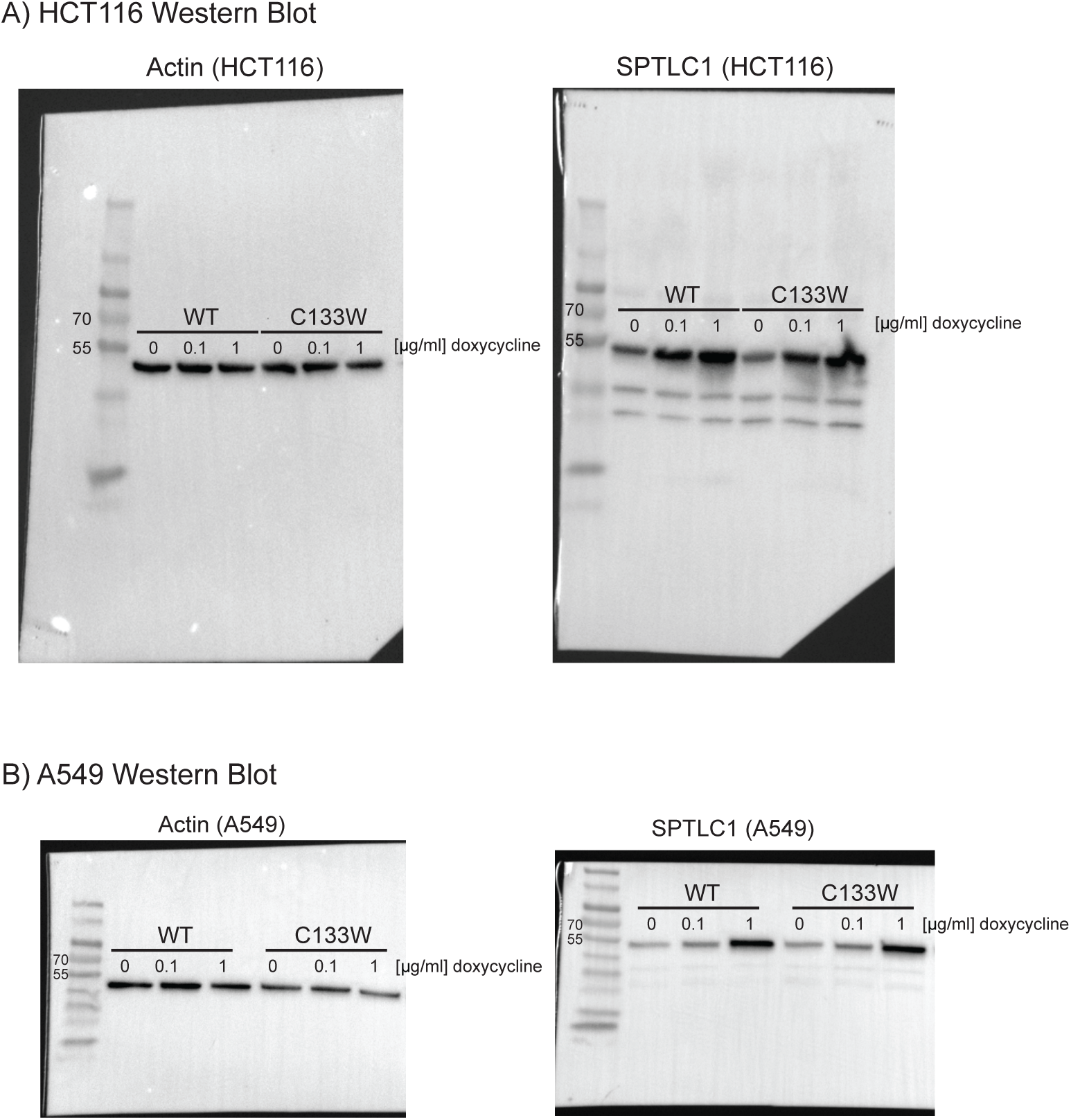
Raw data for immunoblots.

